# Cross-reactive antibody response between SARS-CoV-2 and SARS-CoV infections

**DOI:** 10.1101/2020.03.15.993097

**Authors:** Huibin Lv, Nicholas C. Wu, Owen Tak-Yin Tsang, Meng Yuan, Ranawaka A. P. M. Perera, Wai Shing Leung, Ray T. Y. So, Jacky Man Chun Chan, Garrick K. Yip, Thomas Shiu Hong Chik, Yiquan Wang, Chris Yau Chung Choi, Yihan Lin, Wilson W. Ng, Jincun Zhao, Leo L. M. Poon, J. S. Malik Peiris, Ian A. Wilson, Chris K. P. Mok

## Abstract

The World Health Organization has recently declared the ongoing outbreak of COVID-19, which is caused by a novel coronavirus SARS-CoV-2, as pandemic. There is currently a lack of knowledge in the antibody response elicited from SARS-CoV-2 infection. One major immunological question is concerning the antigenic differences between SARS-CoV-2 and SARS-CoV. We address this question by using plasma from patients infected by SARS-CoV-2 or SARS-CoV, and plasma obtained from infected or immunized mice. Our results show that while cross-reactivity in antibody binding to the spike protein is common, cross-neutralization of the live viruses is rare, indicating the presence of non-neutralizing antibody response to conserved epitopes in the spike. Whether these non-neutralizing antibody responses will lead to antibody-dependent disease enhancement needs to be addressed in the future. Overall, this study not only addresses a fundamental question regarding the antigenicity differences between SARS-CoV-2 and SARS-CoV, but also has important implications in vaccine development.

## Introduction

The emergence of spread of a novel coronavirus SARS-CoV-2 causing severe respiratory disease (COVID-19) has now led to a pandemic with major impact on global health, economy and societal behavior (Coronaviridae Study Group of the International Committee on Taxonomy of, 2020; Poon and Peiris, 2020; Zhu et al., 2020). As of 2020 March 15, over 150,000 confirmed cases of SARS-CoV-2 have been reported with close to 6,000 deaths. Phylogenetic analysis has demonstrated that SARS-CoV-2 and SARS-CoV, a coronavirus that also caused a global outbreak in 2003, are closely related phylogenetically, with genomic nucleotide sequence identity of around 80% (Wu et al., 2020; Zhou et al., 2020). Moreover, it has been shown that both viruses use the angiotensin-converting enzyme 2 (ACE2) as the receptor for cell entry and infection (Letko et al., 2020; Li et al., 2003).

The spike glycoprotein (S) on the surface of coronaviruses is essential for virus entry through binding to the ACE2 receptor and viral fusion with the host cell. The S protein forms a homotrimer in which each protomer is composed of two subunits, S1 and S2 (Figure 1A). Binding between the receptor-binding domain (RBD) in the S1 subunit and the ACE2 receptor triggers a conformational change in the S protein that subsequently initiates membrane fusion events with the host cell. The RBD is also a primary target of the antibody response in humoral immunity and is believed to be the major protective antigen (Chen et al., 2005). The prefusion structure of the S protein of SARS-CoV-2 has been recently determined by cryo-EM (Wrapp et al., 2020), and revealed overall structural similarity to that of SARS-CoV. However, most monoclonal antibodies tested to date that target the RBD of SARS-CoV have failed to bind to the RBD of SARS-CoV-2 (Tian et al., 2020; Wrapp et al., 2020), suggesting that the antigenicity of these two viruses to the RBD is quite distinct. So far, data have not yet been reported from polyclonal human sera from patients to evaluate the antibody response elicited by SARS-CoV-2 infection and to determine whether cross-reactive antibody responses between SARS-CoV-2 and SARS-CoV can be generated. In this study, we examined the antibody responses in 15 patients from Hong Kong who were infected by SARS-CoV-2, and seven by SARS-CoV. Mice infected or immunized with SARS-CoV-2 or SARS-CoV were also used to investigate cross-reactivity of antibody responses between SARS-CoV-2 and SARS-CoV.

**Figure 1.**
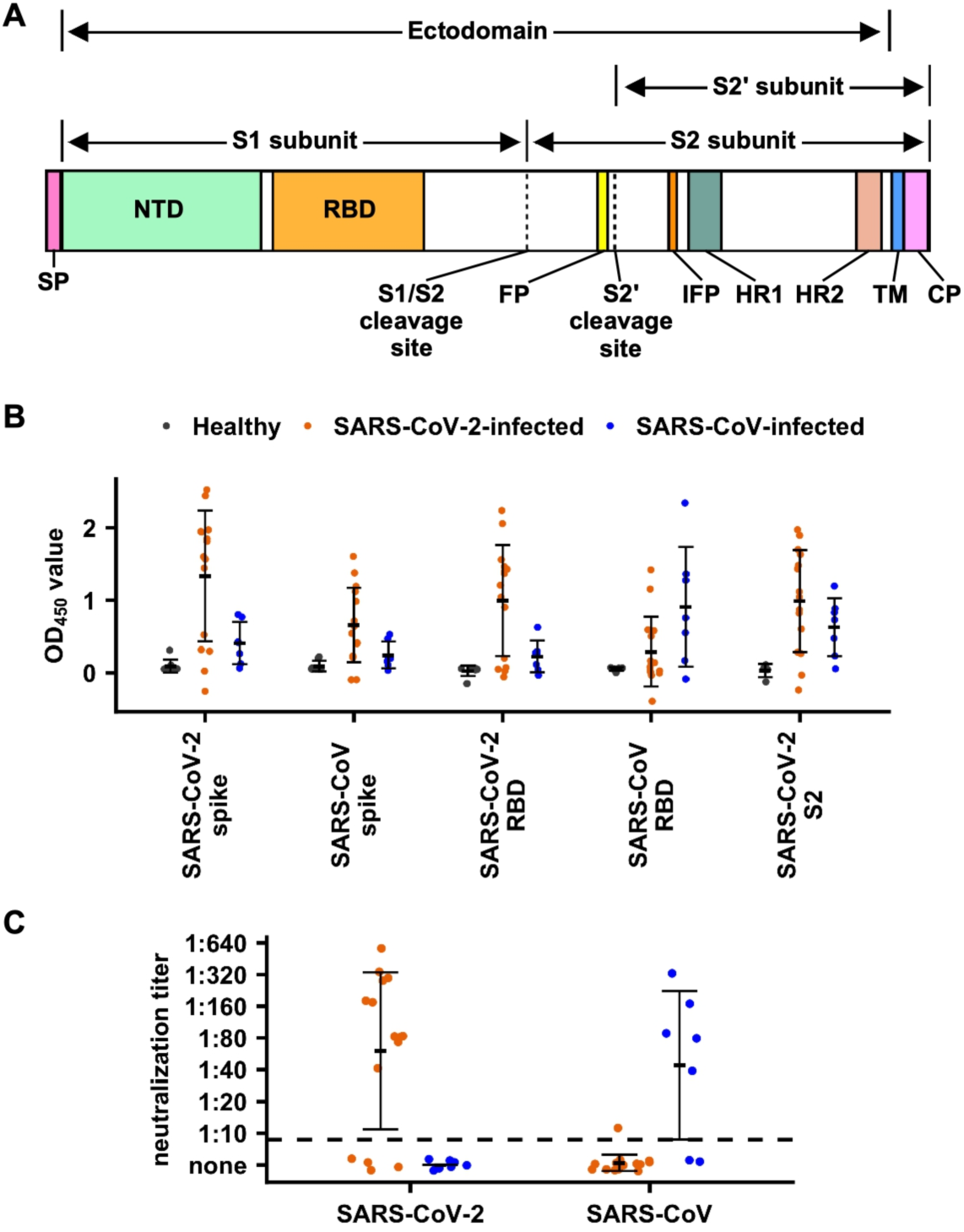
Human serological responses to SARS-CoV-2. **(A)** Schematic diagram of the SARS-CoV-2 spike protein. Locations of secretion signal peptide (SP), N-terminal domain (NTD), receptor-binding domain (RBD), S1/S2 cleavage site, fusion peptide (FP), S2’ cleavage site, internal fusion peptide (IFP), heptad repeat 1 (HR1), heptad repeat 1 (HR2), transmembrane domain (TM), and cytoplasmic domain (CP) are indicated. Regions corresponding to the S1, S2, S2’ subunits, and ectodomain are also indicated. **(B)** Binding of plasma from healthy donors and SARS-CoV-2 infected patients to SARS-CoV-2 spike protein, SARS-CoV-2 RBD protein, SARS-CoV-2 S2 subunit, SARS-CoV spike protein and SARS-CoV RBD protein were measured by ELISA. The mean OD_450_ values calculated after testing each plasma sample in triplicate are shown. **(C)** Neutralization activities of plasma from SARS-CoV-2 infected patients to SARS-CoV-2 and SARS-CoV viruses were measured. Dashed line represents the lower detection limit. Black lines indicate mean +/- standard deviation. **(B-C)** Grey: plasma samples from healthy donors. Orange: plasma samples from SARS-CoV-2-infected patients. Blue: plasma samples from SARS-CoV-infected patients.

## Results

### Patient samples show cross-reactivity in binding

Fifteen heparin anticoagulated plasma samples (from day 2 to 22 post-symptom onset) from SARS-CoV-2 infected patients were analyzed (Table S1). Binding of plasma to the S ectodomain and RBD of both SARS-CoV-2 and SARS-CoV (see Methods) was measured by ELISA (Figure 1B, Figure S1). Plasma samples from healthy donors collected from the Hong Kong Red Cross served as controls. As compared to the plasma from healthy donors, plasma from patients from day 10 post-symptom onward reacted strongly in ELISA binding assays to the S ectodomain (p-value < 2e-16, two-tailed t-test) and RBD (p-value = 2e-13, two-tailed t-test) of SARS-CoV-2. Interestingly, the plasma from SARS-CoV-2-infected patients could also cross-react, although less strongly, with the SARS-CoV S ectodomain (p-value = 8e-06, two-tailed t-test) and the SARS-CoV RBD (p-value = 0.048, two-tailed t-test) (Figure 1B). Nevertheless, only five of the samples from the SARS-CoV-2-infected patients had convincing antibody binding responses to the SARS-CoV RBD. The other plasma reacted more weakly or not at all with the SARS-CoV RBD (Figure 1B). This result indicates that the cross-reactive antibody response to the S protein after SARS-CoV-2 infection mostly targets non-RBD regions. Consistent with that observation, reactivity of the plasma from SARS-CoV-2-infected patients could be detected with the S2 subunit of SARS-CoV-2 (p-value = 2e-4, two-tailed t-test, Figure 1B).

We also analyzed seven heparin anticoagulated convalescent (3-6 months post infection) plasma samples from SARS-CoV infected patients. Similar to that observed in plasma samples from SARS-CoV-2-infected patients, cross-reactivity in binding could be detected (Figure 1B). As compared to the plasma from healthy donors, SARS-CoV-infected patients have significant cross-reactivity in binding to SARS-CoV-2 spike (p-value = 0.03, two-tailed t-test), RBD (p-value = 0.03, two-tailed t-test), and S2 subunit (p-value = 0.007, two-tailed t-test). These results show that cross-reactivity in binding is common between SARS-CoV and SARS-CoV-2 infections in both directions.

### Patient samples show limited cross-neutralization

We next tested the neutralization activity of these plasma samples from SARS-CoV-2-infected patients. Except for four plasma samples that came from patients with less than 12 days post-symptom onset with concomitantly low reactivity to both SARS-CoV-2 S ectodomain and RBD, all other plasma samples could neutralize the SARS-CoV-2 virus with titers ranging from 1:40 to 1:640 (Figure 1C, Table S1). However, only one plasma sample could cross-neutralize SARS-CoV, with very low neutralization activity (1:10). In fact, that cross-neutralizing plasma sample had the strongest reactivity in binding against SARS-CoV S ectodomain among all 15 patient samples, although its binding activity against SARS-CoV RBD is not particularly strong (Table S1).

Similarly, while five of the seven plasma samples from SARS-CoV-convalescent patients could neutralize SARS-CoV with titers ranging from 1:40 to 1:320, none can cross-neutralize SARS-CoV-2 (Figure 1C). These results show that although cross-reactivity in binding is common between plasma from SARS-CoV-2 and SARS-CoV infected patients, cross-neutralization activity is rare.

### Cross-reactivity in mouse infection and vaccination

To further investigate the cross-reactivity of antibody responses to SARS-CoV-2 and SARS-CoV, we analyzed the antibody response of plasma collected from mice infected or immunized with SARS-CoV-2 or SARS-CoV (n = 5 or 6 per experimental and control groups). Plasma from mice with mock immunization with a genetically more distant betacoronavirus coronavirus OC43-CoV, PBS or adjuvant were used as negative controls (Figure 2A-D). As compared to controls, plasma from mice immunized with SARS-CoV-2 virus reacted strongly to its autologous S ectodomain (p-value < 0.002, two-tailed t-test, Figure 2A) and RBD (p-value < 1e-4, two-tailed t-test, Figure 2B). Similarly, plasma from mice immunized with SARS-CoV virus reacted strongly to its autologous S ectodomain (p-value < 2e-7, two-tailed t-test, Figure 2C) and RBD (p-value < 6e-6, two-tailed t-test, Figure 2D). In addition, plasma from mice immunized with SARS-CoV S ectodomain could react to its autologous RBD (p-value < 0.02, two-tailed t-test, Figure 2D). However, while plasma from mice infected with SARS-CoV virus could react with its autologous S ectodomain (p-value < 8e-6, two-tailed t-test, Figure 2C) and RBD (p-value < 2e-5, two-tailed t-test, Figure 2D), the reactivity of plasma from mice infected with SARS-CoV-2 virus to its autologous S ectodomain and RBD could not be observed in this assay (p-value > 0.28, two-tailed t-test, Figure 2A-B). Unlike SARS-CoV virus, which can replicate in wild-type mice (Yang et al., 2004), it has been recently shown that SARS-CoV-2 is only able to replicate in human ACE2-expression transgenic mice but not wild-type mice (Bao et al., 2020), which then can explain the weak immune response from SARS-CoV-2-infected wild-type mice in this study.

**Figure 2.**
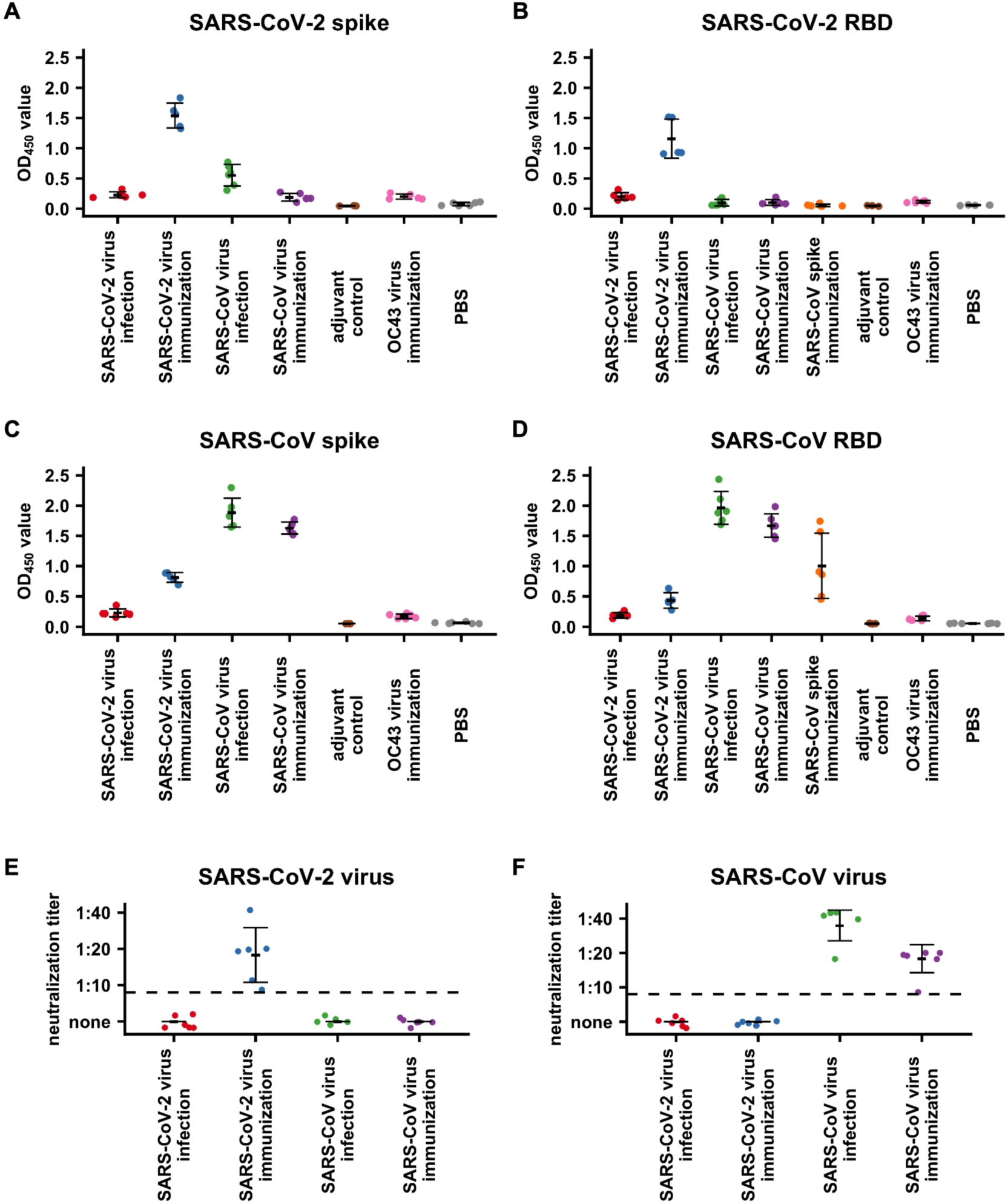
Mouse serological response to SARS-CoV-2 and SARS-CoV. **(A-D)** Binding of plasma from OC43-CoV-immunized mice, SARS-CoV-immunized mice, SARS-CoV-infected mice and mock-immunized mice against **(A)** SARS-CoV-2 spike protein, **(B)** SARS-CoV-2 RBD protein, **(C)** SARS-CoV spike protein and **(D)** SARS-CoV RBD protein were measured by ELISA. Since both SARS-CoV spike protein and SARS-CoV-2 spike contained a C-terminal foldon domain, binding of plasma from mice immunized with SARS-CoV spike protein plasma was not tested against spike proteins from SARS-CoV and SARS-CoV-2. The mean OD_450_ values calculated after testing each plasma sample in triplicate are shown. **(E-F)** Neutralization activities of plasma from mice infected or immunized by SARS-CoV-2 or SARS-CoV to **(E)** SARS-CoV-2 virus or **(F)** SARS-CoV virus were measured. Dashed line represents the lower detection limit. Black lines indicate mean +/- standard deviation.

Interestingly, we observed some cross-reactivity of plasma from SARS-CoV-2-immunized mice to the SARS-CoV S ectodomain (p-value < 4e-5, two-tailed t-test, Figure 2C) and less so to SARS-CoV RBD (p-value < 0.006, two-tailed t-test, Figure 2D), as well as plasma from SARS-CoV-infected mice to the SARS-CoV-2 S ectodomain (p-value < 0.005, two-tailed t-test, Figure 2A). The conclusion that the cross-reactive antibodies mostly target non-RBD regions is supported by the stronger reactivity of the antibody responses from SARS-CoV-2 immunization with the SARS-CoV S ectodomain than to its RBD, and that plasma from SARS-CoV-infected mice did not react at all with SARS-CoV-2 RBD (p-value > 0.5, two-tailed t-test, Figure 2B). Despite the presence of cross-reactivity in binding, cross-neutralization activity was not detected in any of the mouse plasma samples (Figure 2E-F), corroborating with our findings from human patients.

## Discussion

The work here shows that antibody responses in the SARS-CoV-2 infected patient cohort are generated to both S protein and RBD in the majority of the patients. Furthermore, cross-reactivity with SARS-CoV can be detected in these plasma samples as well as in mice studies. These cross-reactive antibody responses mostly target non-RBD regions. Consistently, higher sequence conservation is found between the S2 subunits of SARS-CoV-2 and SARS-CoV (90% amino-acid sequence identity) as compared to that of their RBDs (73% amino-acid sequence identity). Nonetheless, some SARS-CoV-2-infected patients were able to produce cross-reactive antibody responses to SARS-CoV RBD. Consistent with these findings, a human antibody CR3022 that neutralizes SARS-CoV (ter Meulen et al., 2006) has recently been reported to also bind to the RBD of SARS-CoV-2 (Tian et al., 2020).

While cross-reactive antibody binding responses to both SARS-CoV-2 and SARS-CoV S proteins appears to be relatively common in this cohort, cross-neutralizing responses are rare. Only one out of 15 SARS-CoV-2-infected patients was able to generate a cross-neutralizing response to both SARS-CoV-2 and SARS-CoV viruses, and this cross-reactive response was very weak. Therefore, it is possible that only a subset of the cross-reactive binding epitopes is a *bona fide* neutralizing epitope. This notion is also supported by our recent study, which showed that the cross-reactive antibody CR3022 could not neutralize SARS-CoV-2 despite its strong binding (Yuan et al., 2020). Future studies need to investigate whether these non-neutralizing antibody responses can confer *in vivo* protections despite the lack of *in vitro* neutralization activity, which have been observed in some non-neutralizing antibodies to other viruses (Bajic et al., 2019; Bangaru et al., 2019; Bootz et al., 2017; Burke et al., 2018; Dreyfus et al., 2012; Henchal et al., 1988; Lee et al., 2016; Petro et al., 2015; Watanabe et al., 2019). On the contrary, non-neutralizing antibody responses can also lead to antibody-dependent enhancement (ADE) of infection as reported in other coronaviruses (Tseng et al., 2012; Wang et al., 2014; Weiss and Scott, 1981). Whether ADE plays a role in SARS-CoV-2 infection will need to be carefully examined, due to its potential adverse effect in vaccination (Tseng et al., 2012).

SARS-CoV-2 is the third newly emerged coronavirus to cause outbreaks (along with SARS-CoV and MERS-CoV) in the past two decades. Since Coronavirus outbreak are likely to continue to pose global health risks in the future (Menachery et al., 2015; Menachery et al., 2016), the possibility of developing a cross-protective vaccine against multiple coronaviruses has been considered. Identification of cross-protective epitopes on the coronavirus S protein will be important for the development of a more universal coronavirus vaccine analogous to those currently in development for influenza virus. Our findings suggest that such broadly cross-protective epitopes are not common in the human immune repertoire. Moving forward, monoclonal clonal antibodies discovery and characterization will be crucial to the development of a SARS-CoV-2 vaccine in short-term, as well as a cross-protective coronavirus vaccine in long-term.

## Methods

### Recruitment of patients and specimen collections

Patients with RT-PCR confirmed COVID-19 disease at the Infectious Disease Centre of the Princess Margaret Hospital, Hong Kong, were invited to participate in the study after providing informed consent. The study was approved the institutional review board of the Hong Kong West Cluster of the Hospital Authority of Hong Kong (approval number: UW20-169). Specimens of heparinized blood were collected from the patients, and the plasma were separated and stored at -80°C until use. The plasma was heat inactivated at 56°C for 30 minutes before use. The plasma samples from patients with SARS-CoV infection were obtained from the bio-repository of specimens stored from patients following the SARS outbreak in 2003.

### Protein expression and purification

Ectodomain (residues 14-1195) with K968P/V969P mutations and RBD (residues: 306-527) of the SARS-CoV spike (S) protein (GenBank: ABF65836.1), as well as the ectodomain (residues 14-1213) with R682G/R683G/R685G/K986P/V987P mutations and RBD (residues 319-541) of the SARS-CoV-2 spike protein (GenBank: QHD43416.1) were cloned into a customized pFastBac vector (Ekiert et al., 2011). K968P/V969P were stabilizing mutations in the SARS-CoV spike protein (Kirchdoerfer et al., 2018) and the corresponding K986P/V987P mutations in the SARS-CoV-2 spike protein should have the same stabilizing effect due to sequence similarity. R682G/R683G/R685G mutations in the SARS-CoV-2 spike protein were designed to knock-out the furin cleavage site that is a novel addition to this coronavirus compared to related sequences in bats and pangolins (Wong et al., 2020). The spike ectodomain constructs were fused with an N-terminal gp67 signal peptide and a C-terminal BirA biotinylation site, thrombin cleavage site, trimerization domain, and His_6_ tag. The RBD constructs were fused with an N-terminal gp67 signal peptide and a C-terminal His_6_ tag. Recombinant bacmid DNA was generated using the Bac-to-Bac system (Life Technologies). Baculovirus was generated by transfecting purified bacmid DNA into Sf9 cells using FuGENE HD (Promega), and subsequently used to infect suspension cultures of High Five cells (Life Technologies) at an MOI of 5 to 10. Infected High Five cells were incubated at 28 °C with shaking at 110 r.p.m. for 72 h for protein expression. The supernatant was then concentrated using a Centramate cassette (10 kDa MW cutoff for RBD and 30 kDa MW cutoff for spike protein, Pall Corporation). Spike ectodomain and RBD proteins were purified by Ni-NTA (Figure S2), followed by size exclusion chromatography, and then buffer exchanged into PBS. The S2 extracellular domain of SARS-CoV-2 was purchased from Sino Biological, China.

### Mouse immunization

6-8 weeks Balb/c mice were immunized with 10^5^ pfu of SARS-CoV, SARS-CoV-2, HCoV-OC43 or 15 µg of SARS-CoV spike protein in 150 µL PBS together with 50 µL Addavax (MF59-like squalene adjuvant from InvivoGen) through intraperitoneally injection (i.p.). For the control group, Balb/c mice were injected intraperitoneally (i.p.) with 50 µL Addavax plus 150 µL PBS, or 200 µL PBS only. The plasma samples were collected on day 14 post-vaccination using heparin tubes. The experiments were conducted in The University of Hong Kong Biosafety Level 3 (BSL3) facility. This study protocol was carried out in strict accordance with the recommendations and was approved by the Committee on the Use of Live Animals in Teaching and Research of the University of Hong Kong (CULATR 4533-17).

### Mouse infection

6-8 weeks Balb/c mice were anesthetized with Ketamine and Xylazine, and infected intranasally (i.n.) with 10^5^ pfu of SARS-CoV or SARS-CoV-2 diluted in 25 µL PBS. Mouse plasma samples were collected on day 14 post-infection using heparin tubes. The experiments were conducted in the University of Hong Kong Biosafety Level 3 (BSL3) facility.

### ELISA binding assay

A 96-well enzyme-linked immunosorbent assay (ELISA) plate (Nunc MaxiSorp, Thermo Fisher Scientific) was first coated overnight with 100 ng per well of purified recombinant protein in PBS buffer. To substrate the background noise caused by the unspecific binding of antibodies from the samples, serum-specific background noise (SSBN) normalization approach was used (Moritz et al., 2019). In brief, an additional plate was coated overnight with PBS buffer only. The plates coated with either purified recombinant protein or PBS were then blocked with PBS containing 5% non-fat milk powder at room temperature for 2 hours. Each mouse plasma sample was 1:10 diluted and human sample was serially diluted from 1:100 to 1:12800 in PBS. Each sample was then added into the ELISA plates that were coated with purified recombinant protein or PBS buffer respectively for 2-hour incubation at 37°C. After extensive washing with PBS containing 0.1% Tween 20, each well in the plate was further incubated with the HRP-sheep anti-mouse or anti-human secondary antibody (1:5000, GE Healthcare) for 1 hour at 37°C. The ELISA plates were then washed five times with PBS containing 0.1% Tween 20. Subsequently, 50 µL of each solution A and B (R&D Systems) was added into each well. After 15 minutes incubation, the reaction was stopped by adding 50 µL of 2 M H_2_SO_4_ solution and analyzed on a Sunrise (Tecan) absorbance microplate reader at 450 nm wavelength. The normalized results were obtained by the calculating the difference between the OD of the purified recombinant protein-coated well and the PBS-coated well.

### Microneutralization assay

Plasma samples were diluted in serial two-fold dilutions commencing with a dilution of 1:10, and mixed with equal volumes of SARS-CoV or SARS-CoV-2 at a dose of 200 tissue culture infective doses 50% (TCID_50_) determined by Vero and Vero E6 cells respectively. After 1 h of incubation at 37°C, 35 µl of the virus-serum mixture was added in quadruplicate to Vero or Vero E6 cell monolayers in 96-well microtiter plates. After 1 h of adsorption, the virus-serum mixture was removed and replaced with 150ul of virus growth medium in each well. The plates were incubated for 3 days at 37°C in 5% CO_2_ in a humidified incubator. Cytopathic effect was observed at day 3 post-inoculation. The highest plasma dilution protecting 50% of the replicate wells was denoted as the neutralizing antibody titer. A virus back-titration of the input virus was included in each batch of tests.

## Acknowledgements

This work was supported by Calmette and Yersin scholarship (H.L.), NIH K99 AI139445 (N.C.W.), Bill and Melinda Gates Foundation OPP1170236 (I.A.W.), Guangzhou Medical University High-level University Innovation Team Training Program (Guangzhou Medical University released [2017] No.159) (C.K.P.M. and J.S.M.P), the US National Institutes of Health (contract no. HHSN272201400006C) (J.S.M.P), National Natural Science Foundation of China (NSFC)/Research Grants Council (RGC) Joint Research Scheme (N_HKU737/18) (C.K.P.M. and J.S.M.P), and International Cooperation and Exchange of the National Natural Science Foundation of China (Grant No. 8181101118) (J.Z.). We acknowledge the support of the clinicians who facilitated this study including Drs John Yu Hong Chan, Daphne Pui-Lin Lau, and Ying Man Ho and the clinical team at Infectious Diseases Centre, Princess Margaret Hospital, Hospital Authority of Hong Kong and the patients who consented to participate in this investigation.

## Author contributions

H.L., N.C.W., J.S.M.P., I.A.W., C.K.P.M and O.T.Y.T conceived and designed the study. N.C.W. and M.Y. expressed and purified the proteins. H.L., R.T.Y.S., W.W.N., G.K.Y., Y.L., Y.W. and R.A.P.M.P performed the experiments. O.T.Y.T, W.S.L, J.M.C.C, T.S.H.C, and C.Y.C.C. organized patient recruitment, data collection and sampling. H.L., N.C.W., J.Z. L.L.M.P, and C.K.P.M. analyzed the data. H.L., N.C.W., J.S.M.P., I.A.W. and C.K.P.M. wrote the paper and all authors reviewed and edited the paper.

## Competing interests

The authors declare no competing interests.

## References

Bajic, G., Maron, M.J., Adachi, Y., Onodera, T., McCarthy, K.R., McGee, C.E., Sempowski, G.D., Takahashi, Y., Kelsoe, G., Kuraoka, M., et al. (2019). Influenza antigen engineering focuses immune responses to a subdominant but broadly protective viral epitope. Cell Host Microbe 25, 827-835.e826.

Bangaru, S., Lang, S., Schotsaert, M., Vanderven, H.A., Zhu, X., Kose, N., Bombardi, R., Finn, J.A., Kent, S.J., Gilchuk, P., et al. (2019). A site of vulnerability on the influenza virus hemagglutinin head domain trimer interface. Cell 177, 1136-1152.e1118.

Bao, L., Deng, W., Huang, B., Gao, H., Liu, J., Ren, L., Wei, Q., Yu, P., Xu, Y., Qi, F., et al. (2020). The pathogenicity of SARS-CoV-2 in hACE2 transgenic mice. bioRxiv 10.1101/2020.02.07.939389.

Bootz, A., Karbach, A., Spindler, J., Kropff, B., Reuter, N., Sticht, H., Winkler, T.H., Britt, W.J., and Mach, M. (2017). Protective capacity of neutralizing and non-neutralizing antibodies against glycoprotein B of cytomegalovirus. PLoS Pathog 13, e1006601.

Burke, C.W., Froude, J.W., Miethe, S., Hulseweh, B., Hust, M., and Glass, P.J. (2018). Human-like neutralizing antibodies protect mice from aerosol exposure with western equine encephalitis virus. Viruses 10, 147.

Chen, Z., Zhang, L., Qin, C., Ba, L., Yi, C.E., Zhang, F., Wei, Q., He, T., Yu, W., Yu, J., et al. (2005). Recombinant modified vaccinia virus Ankara expressing the spike glycoprotein of severe acute respiratory syndrome coronavirus induces protective neutralizing antibodies primarily targeting the receptor binding region. J Virol 79, 2678–2688.

Coronaviridae Study Group of the International Committee on Taxonomy of, V. (2020). The species Severe acute respiratory syndrome-related coronavirus: classifying 2019-nCoV and naming it SARS-CoV-2. Nat Microbiol 10.1038/s41564-020-0695-z.

Dreyfus, C., Laursen, N.S., Kwaks, T., Zuijdgeest, D., Khayat, R., Ekiert, D.C., Lee, J.H., Metlagel, Z., Bujny, M.V., Jongeneelen, M., et al. (2012). Highly conserved protective epitopes on influenza B viruses. Science 337, 1343–1348.

Ekiert, D.C., Friesen, R.H., Bhabha, G., Kwaks, T., Jongeneelen, M., Yu, W., Ophorst, C., Cox, F., Korse, H.J., Brandenburg, B., et al. (2011). A highly conserved neutralizing epitope on group 2 influenza A viruses. Science 333, 843–850.

Henchal, E.A., Henchal, L.S., and Schlesinger, J.J. (1988). Synergistic interactions of anti-NS1 monoclonal antibodies protect passively immunized mice from lethal challenge with dengue 2 virus. J Gen Virol 69 (Pt 8), 2101–2107.

Kirchdoerfer, R.N., Wang, N., Pallesen, J., Wrapp, D., Turner, H.L., Cottrell, C.A., Corbett, K.S., Graham, B.S., McLellan, J.S., and Ward, A.B. (2018). Stabilized coronavirus spikes are resistant to conformational changes induced by receptor recognition or proteolysis. Sci Rep 8, 15701.

Lee, J., Boutz, D.R., Chromikova, V., Joyce, M.G., Vollmers, C., Leung, K., Horton, A.P., DeKosky, B.J., Lee, C.H., Lavinder, J.J., et al. (2016). Molecular-level analysis of the serum antibody repertoire in young adults before and after seasonal influenza vaccination. Nat Med 22, 1456–1464.

Letko, M., Marzi, A., and Munster, V. (2020). Functional assessment of cell entry and receptor usage for SARS-CoV-2 and other lineage B betacoronaviruses. Nat Microbiol 10.1038/s41564-020-0688-y.

Li, W., Moore, M.J., Vasilieva, N., Sui, J., Wong, S.K., Berne, M.A., Somasundaran, M., Sullivan, J.L., Luzuriaga, K., Greenough, T.C., et al. (2003). Angiotensin-converting enzyme 2 is a functional receptor for the SARS coronavirus. Nature 426, 450–454.

Menachery, V.D., Yount, B.L., Jr., Debbink, K., Agnihothram, S., Gralinski, L.E., Plante, J.A., Graham, R.L., Scobey, T., Ge, X.Y., Donaldson, E.F., et al. (2015). A SARS-like cluster of circulating bat coronaviruses shows potential for human emergence. Nat Med 21, 1508–1513.

Menachery, V.D., Yount, B.L., Jr., Sims, A.C., Debbink, K., Agnihothram, S.S., Gralinski, L.E., Graham, R.L., Scobey, T., Plante, J.A., Royal, S.R., et al. (2016). SARS-like WIV1-CoV poised for human emergence. Proc Natl Acad Sci U S A 113, 3048–3053.

Moritz, C.P., Tholance, Y., Lassabliere, F., Camdessanche, J.P., and Antoine, J.C. (2019). Reducing the risk of misdiagnosis of indirect ELISA by normalizing serum-specific background noise: The example of detecting anti-FGFR3 autoantibodies. J Immunol Methods 466, 52–56.

Petro, C., Gonzalez, P.A., Cheshenko, N., Jandl, T., Khajoueinejad, N., Benard, A., Sengupta, M., Herold, B.C., and Jacobs, W.R. (2015). Herpes simplex type 2 virus deleted in glycoprotein D protects against vaginal, skin and neural disease. eLife 4, e06054.

Poon, L.L.M., and Peiris, M. (2020). Emergence of a novel human coronavirus threatening human health. Nat Med 10.1038/s41591-020-0796-5.

ter Meulen, J., van den Brink, E.N., Poon, L.L., Marissen, W.E., Leung, C.S., Cox, F., Cheung, C.Y., Bakker, A.Q., Bogaards, J.A., van Deventer, E., et al. (2006). Human monoclonal antibody combination against SARS coronavirus: synergy and coverage of escape mutants. PLoS Med 3, e237.

Tian, X., Li, C., Huang, A., Xia, S., Lu, S., Shi, Z., Lu, L., Jiang, S., Yang, Z., Wu, Y., et al. (2020). Potent binding of 2019 novel coronavirus spike protein by a SARS coronavirus-specific human monoclonal antibody. Emerg Microbes Infect 9, 382–385.

Tseng, C.T., Sbrana, E., Iwata-Yoshikawa, N., Newman, P.C., Garron, T., Atmar, R.L., Peters, C.J., and Couch, R.B. (2012). Immunization with SARS coronavirus vaccines leads to pulmonary immunopathology on challenge with the SARS virus. PLoS One 7, e35421.

Wang, S.F., Tseng, S.P., Yen, C.H., Yang, J.Y., Tsao, C.H., Shen, C.W., Chen, K.H., Liu, F.T., Liu, W.T., Chen, Y.M., et al. (2014). Antibody-dependent SARS coronavirus infection is mediated by antibodies against spike proteins. Biochem Biophys Res Commun 451, 208–214.

Watanabe, A., McCarthy, K.R., Kuraoka, M., Schmidt, A.G., Adachi, Y., Onodera, T., Tonouchi, K., Caradonna, T.M., Bajic, G., Song, S., et al. (2019). Antibodies to a conserved influenza head interface epitope protect by an IgG subtype-dependent mechanism. Cell 177, 1124-1135.e1116.

Weiss, R.C., and Scott, F.W. (1981). Antibody-mediated enhancement of disease in feline infectious peritonitis: comparisons with dengue hemorrhagic fever. Comp Immunol Microbiol Infect Dis 4, 175–189.

Wong, M.C., Javornik Cregeen, S.J., Ajami, N.J., and Petrosino, J.F. (2020). Evidence of recombination in coronaviruses implicating pangolin origins of nCoV-2019. bioRxiv 10.1101/2020.02.07.939207.

Wrapp, D., Wang, N., Corbett, K.S., Goldsmith, J.A., Hsieh, C.L., Abiona, O., Graham, B.S., and McLellan, J.S. (2020). Cryo-EM structure of the 2019-nCoV spike in the prefusion conformation. Science 10.1126/science.abb2507.

Wu, F., Zhao, S., Yu, B., Chen, Y.M., Wang, W., Song, Z.G., Hu, Y., Tao, Z.W., Tian, J.H., Pei, Y.Y., et al. (2020). A new coronavirus associated with human respiratory disease in China. Nature 10.1038/s41586-020-2008-3.

Yang, Z.Y., Kong, W.P., Huang, Y., Roberts, A., Murphy, B.R., Subbarao, K., and Nabel, G.J. (2004). A DNA vaccine induces SARS coronavirus neutralization and protective immunity in mice. Nature 428, 561–564.

Yuan, M., Wu, N.C., Zhu, X., Lee, C.-C.D., So, R.T.Y., Lv, H., Mok, C.K.P., and Wilson, I.A. (2020). A highly conserved cryptic epitope in the receptor-binding domains of SARS-CoV-2 and SARS-CoV. bioRxiv 10.1101/2020.03.13.991570.

Zhou, P., Yang, X.L., Wang, X.G., Hu, B., Zhang, L., Zhang, W., Si, H.R., Zhu, Y., Li, B., Huang, C.L., et al. (2020). A pneumonia outbreak associated with a new coronavirus of probable bat origin. Nature 10.1038/s41586-020-2012-7.

Zhu, N., Zhang, D., Wang, W., Li, X., Yang, B., Song, J., Zhao, X., Huang, B., Shi, W., Lu, R., et al. (2020). A novel coronavirus from patients with pneumonia in China,2019. N Engl J Med 382, 727–733.

