## Supplementary Materials for "Cross-reactive antibody response between SARS-CoV-2 and SARS-CoV infections"

### Supplementary Table 1. Patient information, ELISA data, and neutralization data.

| | | | | ELISA ($OD_{450}$ value at 1:100 dilution) | | | | MN titer | |
|---|---|---|---|---|---|---|---|---|---|
| Patient ID | Gender | Age | day post-symptom onset | SARS-CoV-2 spike | SARS-CoV spike | SARS-CoV-2 RBD | SARS-CoV RBD | SARS-CoV-2 | SARS-CoV |
| #1 | M | 56 | 6 | 0.30 | 0.21 | 0.03 | 0.01 | <1:10 | <1:10 |
| #2 | F | 62 | 4 | 0.32 | 0.23 | 0.07 | 0.03 | <1:10 | <1:10 |
| #3 | F | 62 | 22 | 1.97 | 0.66 | 1.46 | 0.12 | 1:160 | <1:10 |
| #4 | M | 63 | 22 | 1.95 | 0.98 | 1.56 | 0.58 | 1:80 | <1:10 |
| #5 | M | 47 | 8 | 0.52 | 0.40 | 0.20 | 0.03 | 1:80 | <1:10 |
| #6 | M | 64 | 19 | 1.81 | 0.54 | 1.40 | 0.50 | 1:320 | <1:10 |
| #7 | F | 73 | 18 | 1.60 | 0.71 | 0.90 | 0.17 | 1:80 | <1:10 |
| #8 | M | 72 | 18 | 1.94 | 1.19 | 1.21 | 0.14 | 1:80 | <1:10 |
| #9 | F | 37 | 2 | -0.25 | -0.10 | -0.05 | -0.39 | <1:10 | <1:10 |
| #10 | F | 72 | 10 | 1.44 | 0.41 | 1.04 | -0.03 | 1:40 | <1:10 |
| #11 | M | 60 | 22 | 2.53 | 1.38 | 2.24 | 1.15 | 1:160 | <1:10 |
| #12 | M | 56 | 13 | 1.57 | 0.64 | 1.37 | 0.00 | 1:320 | <1:10 |
| #13 | F | 55 | 11 | 0.03 | -0.09 | 0.05 | 0.08 | <1:10 | <1:10 |
| #14 | F | 63 | 19 | 2.44 | 1.12 | 2.06 | 1.42 | 1:640 | <1:10 |
| #15 | M | 37 | 14 | 1.85 | 1.60 | 1.43 | 0.59 | 1:320 | 1:10 |

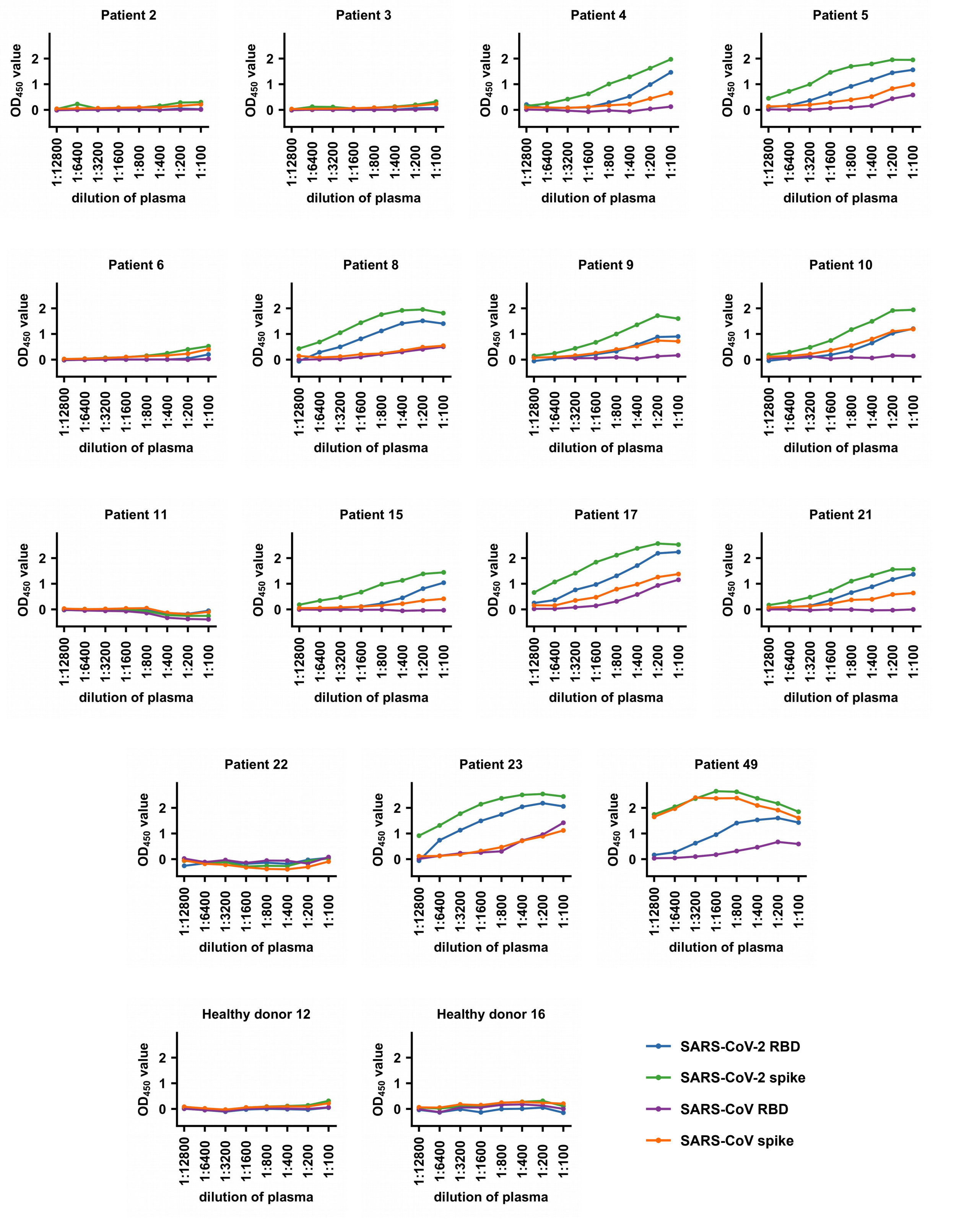


**Supplementary Figure 1. ELISA of patient samples at different dilutions.** Binding of different dilutions of plasma samples from patients and healthy donors to spike and RBD from SARS-CoV-2 and SARS-CoV was measured by ELISA. The mean $OD_{450}$ values calculated after testing each plasma sample in duplicate are shown.

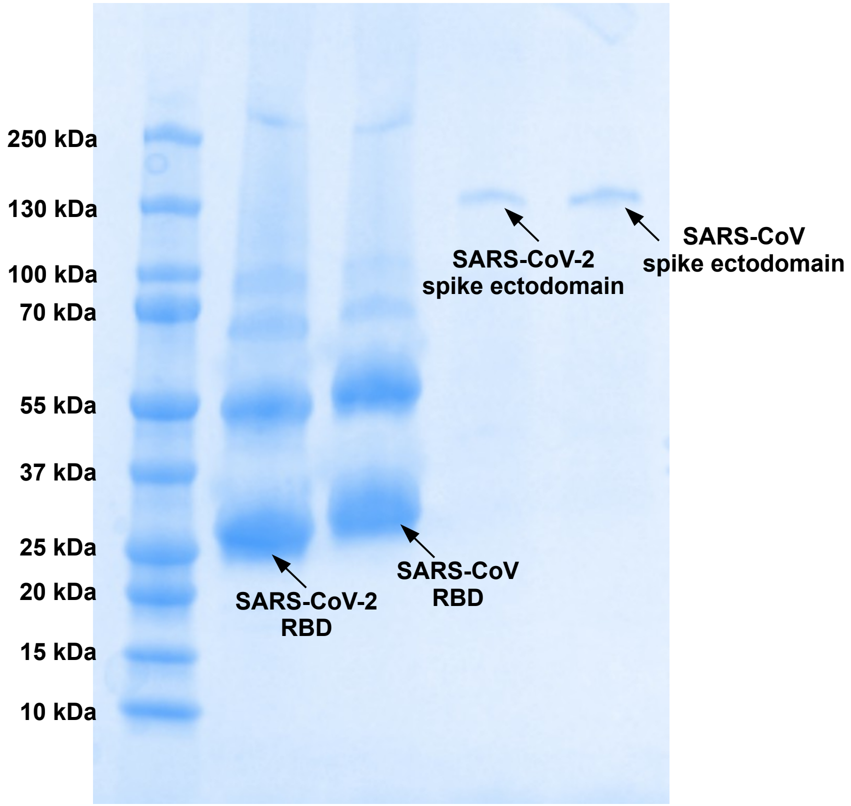


**Supplementary Figure 2. Protein expression.** Proteins purified by Ni-NTA beads from insect cell expression supernatant were analyzed by coomassie staining. The proteins were further purified by size exclusion chromatography before used.
